# HIV Nef amplifies mechanical heterogeneity to promote immune evasion

**DOI:** 10.64898/2025.12.24.696400

**Authors:** Louise Leyre, Farah Mustapha, Alberto Herrera, Esther Lee, Emily Huntsman, Paul Zumbo, Jared Weiler, Parul Sinha, Ethan Naing, Connor Smith, Colin Kovacs, Micheal Galiano, Nada Wahman, Doron Betel, Kiera L. Clayton, Morgan Huse, R. Brad Jones

**Affiliations:** Infectious Disease Division, Weill Cornell Medicine, New York, NY, USA; Immunology and Microbial Pathogenesis Program, Weill Cornell Graduate School of Medical Sciences, New York, NY, USA; Immunology Program, Memorial Sloan Kettering Cancer Center, New York, NY, USA; Applied Bioinformatics Core, Weill Cornell Medicine, New York, NY, USA; Molecular Cytology Core Facility, Memorial Sloan Kettering Cancer Center, New York, NY, USA; Maple Leaf Medical Clinic and Division of Infectious Diseases, Department of Medicine, University of Toronto, Toronto, Canada; Department of Pathology, University of Massachusetts Chan Medical School, Worcester, MA, USA

## Abstract

Intracellular pathogens must evade cytotoxic immunity to establish persistent infection. Although immune escape is typically viewed through a biochemical lens, the ability of certain pathogens to alter the mechanical properties of infected cells suggests that biophysical mechanisms may also contribute to the process. Here, we show that a subset of CD4^+^ T cells infected with the human immunodeficiency virus (HIV) resist elimination through a soft phenotype that inhibits killing by mechanosensitive cytotoxic T lymphocytes (CTLs). This phenotype arises from the combined effects of the HIV virulence factor Nef, which remodels the actin cytoskeleton, and intrinsic heterogeneity in the basal cytoskeletal properties of infected T cells. Pharmacological or genetic perturbations that reverse Nef signaling to the cytoskeleton or that stiffen the filamentous-actin cortex sensitize infected cells to CTL-mediated lysis. Taken together, these findings define a novel, biophysical paradigm of immune evasion with implications for HIV cure strategies.

## Introduction

The ability of the immune system to detect and eliminate infected cells has driven the evolution of evasion mechanisms that enable intracellular pathogens to resist both cellular and humoral immunity^1,2^. Deciphering these mechanisms is critical for understanding disease pathogenesis and for developing effective interventions. HIV provides a paradigmatic example: in untreated infection, the virus circumvents cellular immune pressure through rapid mutational escape within CTL-targeted epitopes^3,4^, Nef-mediated downregulation of MHC-I^5^, and progressive exhaustion of antiviral effector cells^6–9^. Despite vigorous immune responses, the virus ultimately prevails, culminating in progressive immunodeficiency and AIDS.

Antiretroviral therapy (ART) durably suppresses HIV replication but does not eliminate the rebound-competent reservoir, composed predominantly of CD4^+^ T cells harboring intact proviruses, that persists for life and drives rapid viral rebound upon treatment interruption^10–13^. The mechanisms enabling these long-lived infected cells to survive immune attack remain incompletely understood. While latency, which renders infected cells antigenically silent, has long been viewed as the principal barrier to clearance, latency reversal clinical trials have revealed that viral reactivation alone does not ensure CTL elimination^14–16^. These outcomes have coincided with a growing recognition that HIV latency is not absolute; low-level HIV RNA and protein expression persist even under ART, and both have been shown to correlate with the magnitude and functionality of HIV-specific CTLs^17–20^. These observations suggest that, beyond latency, additional mechanisms of immune evasion actively protect infected cells from CTL-mediated elimination.

In untreated infection, immune escape proceeds through the selection of viral variants bearing escape mutations within CTL-targeted epitopes. By contrast, ART arrests viral replication and therefore halts viral evolution. Yet, CD4⁺ T cells harboring intact proviruses continue to undergo clonal expansion during ART, raising the possibility that selective pressures act at the cellular rather than the viral level, favoring the persistence of infected cells intrinsically resistant to CTL attack^21,22^. In prior work, we identified overexpression of the pro-survival factor BCL-2 as one such mechanism^23,24^. Pharmacologic inhibition of BCL-2 sensitized reservoir-harboring cells to CTL-mediated killing *ex vivo*, but this approach only partially depleted the reservoir and incurred substantial bystander toxicity^23,25,26^. This motivated a search for additional, mechanistically distinct and therapeutically tractable pathways of resistance beyond apoptotic regulation.

HIV infection is known to remodel filamentous (F)-actin, producing characteristic changes in cytoskeletal configuration and overall cell morphology^27,28^. These remodeling events, while essential for viral entry, intracellular trafficking, and budding of virus, also have the potential to impact immune recognition^27,28^. This is because cellular cytotoxicity is a fundamentally mechanical process. To kill, cytotoxic lymphocytes engage their target through a structurally stereotyped immune synapse, into which they secrete the pore-forming protein perforin along with several granzyme proteases^29–31^. The immune synapse is a highly dynamic contact, imparting nanonewton-scale forces against the target surface that promote lymphocyte activation, guide cytotoxic secretion, and potentiate perforin pore formation^32,33^. Given the importance of interfacial mechanics for these processes, the physical properties of target cells could influence their vulnerability to CTL attack. Indeed, tumor cells can evade CTLs and natural killer (NK) cells by adopting mechanically pliable states that attenuate the formation of potent immune synapses and thereby limit cytotoxic effector function^34–36^. Whether infectious agents employ analogous escape mechanisms is not known.

Here, we identify a previously unrecognized form of HIV immune evasion that operates through biophysical, rather than purely biochemical, mechanisms. We show that HIV infection alters the mechanical properties of CD4⁺ T cells in a manner that limits their susceptibility to CTL-mediated killing. This resistance arises partly from intrinsic variability within the host CD4^+^ T cell population but is strongly amplified by the HIV virulence factor Nef and can be reversed by pharmacologic restoration of actin dynamics. These findings define a novel, mechanical axis of immune evasion in HIV-infected cells and point to new therapeutic opportunities for sensitizing viral reservoirs to immune clearance.

## Results

### A subset of HIV-expressing CD4^+^ T cells resists CTL killing

Prior transcriptomic efforts to understand the resistance of HIV-infected cells to CTLs were limited by their inability to distinguish intrinsic resistance programs from cellular responses to infection and the inflammatory milieu^23,37^. To overcome these deficiencies, we established a dual-virus co-culture model that enables the direct and internally controlled comparison of targeted versus untargeted HIV-infected cells within the same environment. Naïve CD4⁺ T cells from an HIV-negative donor, matched for the HLA restriction of HIV-specific CD8⁺ CTL clones recognizing either the Gag TW10 (TSTLQEQIGW; HLA-B57/B58) or SL9 (SLYNTVATL; HLA-A02) epitopes, were activated into Tcm-like memory cells (Figure S1A), and infected with either wild-type HIV-JRCSF (HIV WT) or an epitope escape variant harboring the T242N mutation in the Gag TW10 epitope (HIV escape), which abrogates recognition by TW10-specific CTLs (Figure S1B, C). These two infected populations were mixed and co-cultured at a 1:1:1 effector-to-target (E:T) ratio with TW10-specific CTLs. Because T242N-bearing targets are invisible to TW10 CTLs, HIV escape-infected cells served as stringent internal “bystanders,” experiencing identical infection and cytokine environments (e.g., IFN-γ exposure) but without antigen-specific killing (Figure 1A).

**Figure 1:**
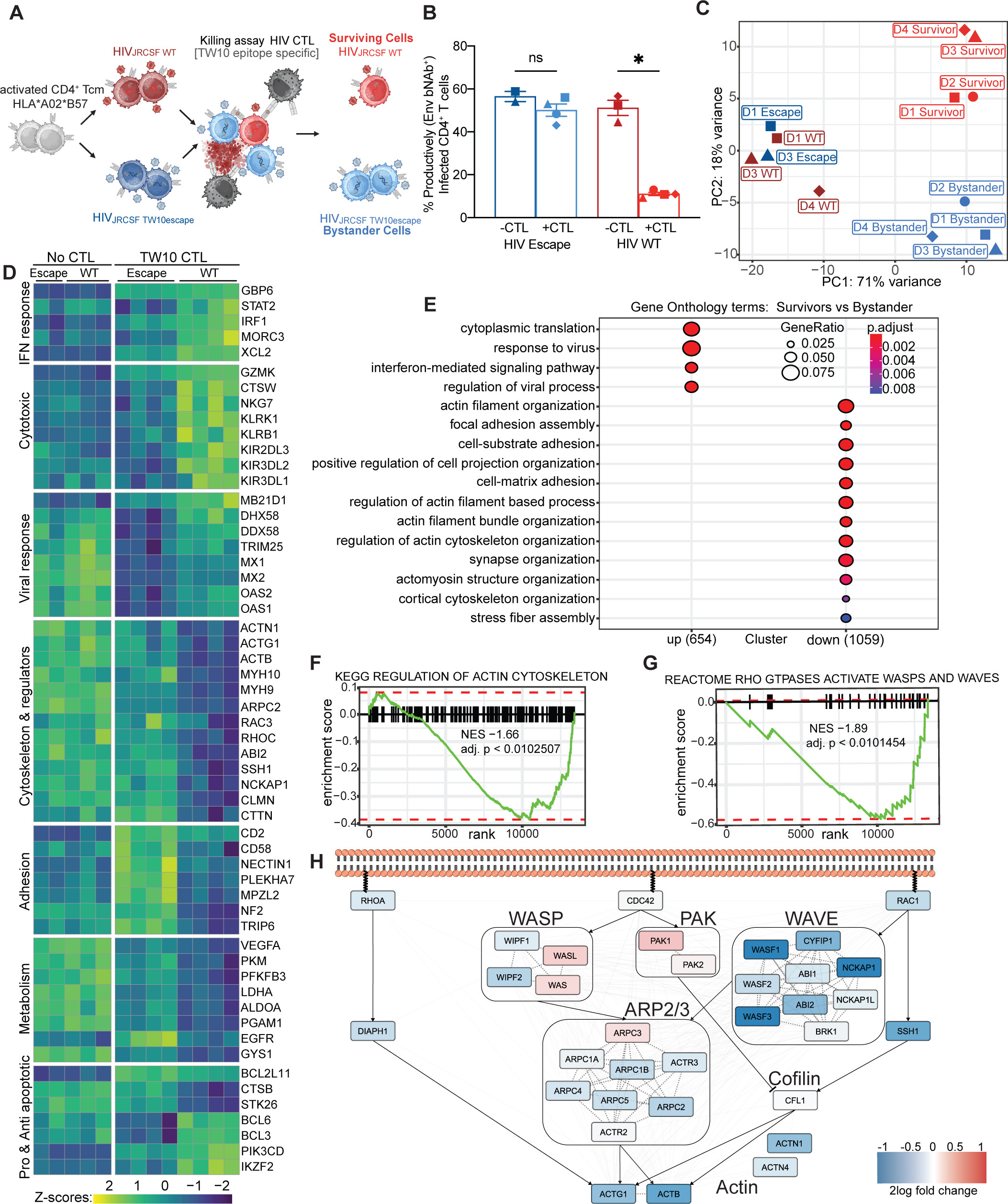
HIV-infected CD4^+^ T cells that survive CTL-mediated killing display a distinct transcriptional profile and cytoskeletal program. (A) Schematic of killing assay in which HIV-Gag-TW10-specific CTLs were co-cultured for 16 h with HIV-WT (red) or HIV-escape (blue) infected cells. (B) Percentage of productively infected (Env bNAb+) CD4+ T cells with or without CTL exposure, showing elimination of HIV-WT but not HIV-escape targets. (C) Principal component analysis (PCA) of RNA-seq profiles of HIV-WT and HIV-escape infected cells without CTL co-culture (dark red and dark blue) and of surviving HIV-WT cells (red) and HIV-escape bystander cells (blue) after CTL exposure (D1 and D2 = donor 1 and donor 2). (D) Heatmap of differentially expressed genes (DEGs) in HIV-WT and HIV-escape infected cells with or without CTLs (row-normalized log counts per million; Wald test-*P*-adj < 0.001). (E) Gene Ontology (GO) enrichment of survivor versus bystander transcriptomes reveals suppression of actin-related processes (GeneRatio >20/X. *P*-adj <0.01). (F and G) Gene set enrichment analysis (GSEA) identifies down-regulation of KEGG *Regulation of Actin Cytoskeleton* and Reactome *Rho GTPases Activate WASPs and WAVEs* pathways in survivor cells. NES = normalized enrichment score. (H) Network map of actin-remodeling genes showing broad repression of cytoskeletal machinery in CTL-resistant survivors. Blue and red denote down- and up-regulation in survivor cells, respectively.

After overnight co-culture with TW10 CTLs, HIV WT-infected cells were substantially depleted (10.6-fold reduction, 0.00138 p value), while HIV escape-infected cells were largely unaffected (1.0-fold reduction, 0.4798 p value) (Figure 1B, S1D). Notably, a fraction of HIV WT-infected cells persisted. To determine whether these survivors were intrinsically resistant to CTL killing, we replaced TW10 CTLs with SL9-specific CTLs, which recognize both WT and escape viruses (Figure S1G). SL9 CTLs strongly depleted the HIV escape-infected targets (8.7-fold reduction) without further reducing the HIV WT-infected pool (Figure S1H), providing direct evidence that surviving WT HIV-infected T cells were intrinsically resistant to CTLs under conditions of effective antigen recognition. This platform thus allowed us to isolate transcriptional and phenotypic features specifically associated with CTL-targeted cells, and to define intrinsic programs linked to resistance.

### CTL-resistant HIV-infected CD4^+^ T cells downregulate cytoskeletal remodeling programs

We next profiled the transcriptional signature of the survivor population identified above. Survivor (HIV WT-infected) and bystander (HIV escape-infected) CD4^+^ T cells from TW10 CTL co-cultures were live-sorted based on surface HIV Env expression (Figure S1E-F), and subjected to bulk RNA-seq. In the absence of CTLs, HIV WT- and escape-infected cells exhibited nearly identical transcriptomes (35 differentially expressed genes), indicating that the T242N escape mutation did not substantially alter the impact of HIV on host gene expression. Following CTL co-culture, however, survivors displayed 1,713 differentially expressed genes (DEGs) (*P*-adj < 0.05), 1,059 of which were downregulated relative to bystanders (Figure 1C-D, S2A). Transcriptional features previously associated with CTL resistance were recapitulated here, including upregulation of transcripts involved in interferon response, translation and cytotoxicity, as well as downregulation of metabolic transcripts related to hypoxia, glycolysis, and apoptosis^38^ (Figure 1D, S2B-C). While BCL-2 itself showed only a modest increase in survivors versus bystanders (0.21-fold, 0.11 p value), its pro-apoptotic BH3-only counterpart BIM was significantly under-expressed (-0.45-fold, 0.02 p value), consistent with enhanced mitochondrial resilience and reduced apoptotic priming (Figure S2D, E). Hence, the two-virus co-culture system faithfully reproduces the metabolic and survival-associated signatures of CTL-resistant cells^38^, validating its capacity to identify and dissect the mechanisms underlying this resistance.

Among the most coherent transcriptional shifts in survivors was a broad suppression of cytoskeleton remodeling pathways, consistent with attenuated actin dynamics. Gene Ontology (GO) analysis revealed marked downregulation of genes associated with actin filament organization, stress fiber formation, adhesion assembly, and other pathways governing the F-actin cytoskeleton (Figure 1E). Several of the most suppressed gene sets converged on cytoskeletal remodeling, a process known to govern immune synapse stability and target cell susceptibility to CTL-mediated killing (Figure 1E, D)^39,40^. Conversely, upregulated genes were enriched for GO pathways linked to antiviral responses and interferon signaling (Figure 1E and S2B), potentially reflecting engagement of cytokine-independent signaling within the immunological synapse or the polarized delivery of cytokines directly into the CTL-target intercellular space^41–43^. Gene set enrichment analysis corroborated these findings, revealing significant downregulation of the KEGG “Regulation of Actin Cytoskeleton” and REACTOME “Rho GTPases Activate WASPs and WAVEs” modules (Figure 1F-H). Together, these data define a transcriptional state marked by repression of cytoskeletal remodeling as a hallmark of survivor cells, suggesting either active adaptation to CTL engagement or preferential survival of a pre-existing mechanically resilient subset.

### CTL-resistant CD4^+^ T cells exhibit reduced cortical and membrane tension

Given the central role of F-actin in governing cellular mechanics, we hypothesized that the reduced expression of cytoskeletal genes in survivor cells reflects the selective persistence of infected cells with intrinsically lower cortical stiffness and altered membrane mechanics. To test this, we performed single-cell measurements of cortical stiffness (Young’s modulus) by atomic force microscopy (AFM) (Figure 2A). We also measured plasma membrane tension by using an optical trap to pull a thin tether of membrane from the cell surface and then quantifying the restoring force applied by the tether on the trap (Figure 2D). Prior to CTL exposure, both WT and escape-infected cells exhibited a lower cortical stiffness relative to uninfected controls (Figure 2B, C), indicating that HIV infection alone partially softens the cell cortex. In contrast, membrane tension was comparable across all groups in the absence of CTL pressure (Figure 2E, F). Following CTL coculture, however, survivor cells displayed a pronounced decrease in both cortical stiffness and membrane tension. Importantly, the mechanical profiles of bystander and uninfected cells were unaltered by the presence of CTLs, indicating that softening is not a generalized consequence of CTL exposure but reflects the selective survival of a mechanically distinct subset of HIV-infected cells. Together, these findings support a model in which CTLs preferentially eliminate stiffer, higher-tension targets, allowing softer, mechanically compliant infected cells to persist.

**Figure 2:**
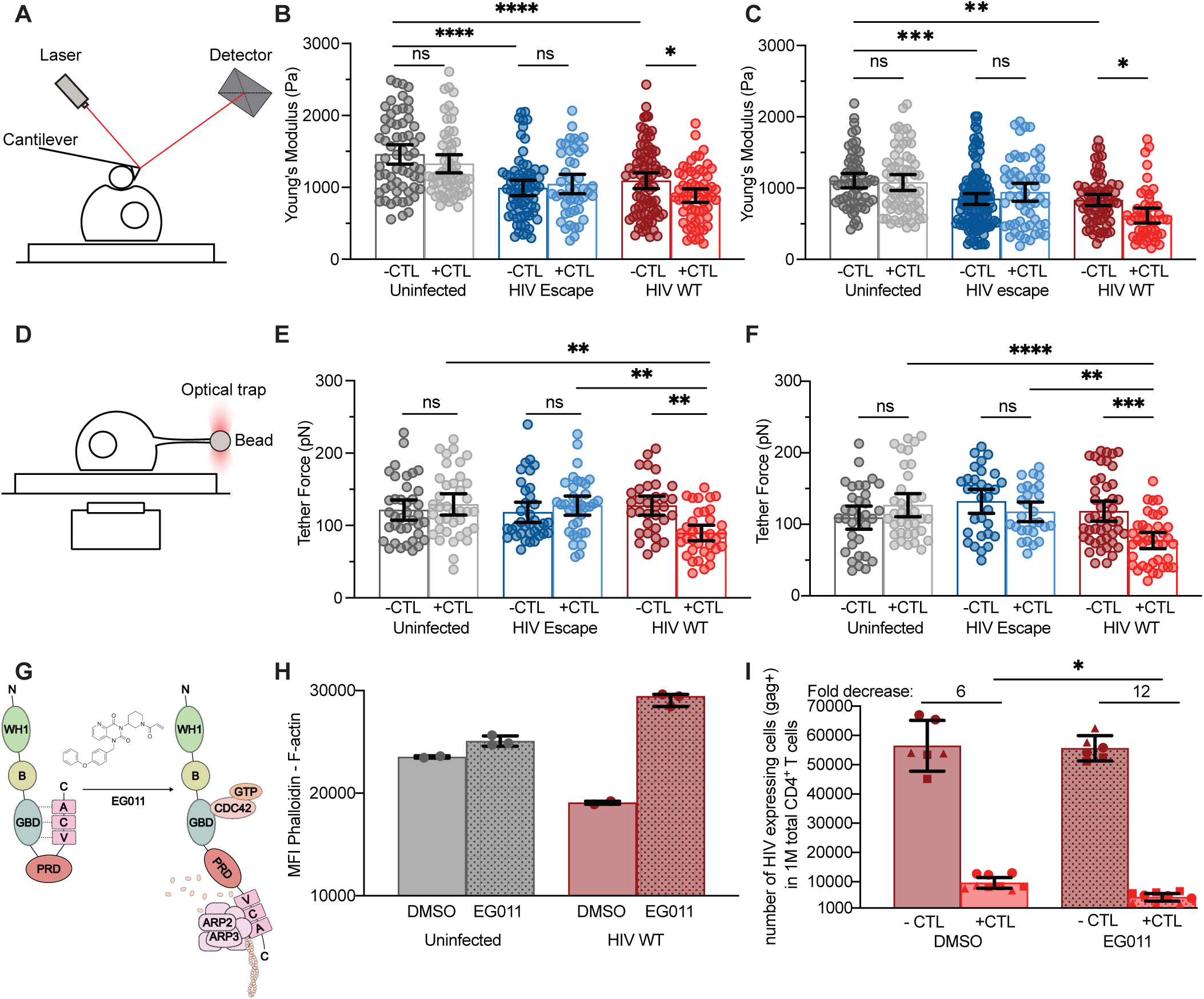
HIV infected CD4^+^ T cells undergo mechanical softening, a phenotype that becomes more pronounced in cells resistant to CTL-mediated killing. (A) Schematic of AFM setup used to measure cell cortical stiffness (Young’s modulus, Pa) in fixed CD4^+^ T cells. (B and C) Cortical stiffness of uninfected (gray), HIV escape-infected (blue), and HIV WT-infected (red) CD4^+^ T cells with and without CTL exposure, shown for two donors. Each point represents an individual cell; mean ± 95% CI from two independent experiments per condition (one-way ANOVA, p<0.0001). (D) Schematic of the optical trap assay used to measure tether force (piconewton, pN) in live cells. (E and F) Tether force measurements for uninfected (gray), HIV escape-infected (blue) and HIV WT-infected (red) CD4^+^ T cells with and without CTL exposure, shown for two donors. Each point represents an individual cell; mean ± 95% CI from two independent experiments per condition (one-way ANOVA, p<0.0001). (G) Schematic diagram showing WASP activation by the allosteric activator EG011. (H) F-actin abundance measured by phalloidin staining and flow cytometry in uninfected and HIV-WT cells treated for 24 h with EG011 (1.5μM) or vehicle (DMSO). (I) CTL killing assay of HIV-WT-infected cells pretreated for 6h with EG011 (1.5μM) or vehicle (DMSO), followed by co-culture with or without TW10 CTLs in the continued presence of drug for 16 h. The number of Gag^+^ cells per 1M CD4^+^ T cells is shown across three donors (circles, squares, triangles; one-way ANOVA: p<0.0001; EG011 + CTL vs. DMSO +CTL, p=0.02. Mean ± 95% CI shown).

To determine whether restoring actin dynamics could reverse this resistant phenotype, we treated HIV WT-infected cells with EG011, a small-molecule activator of the F-actin nucleation protein WASP, which promotes Arp2/3-mediated actin polymerization (Figure 2G)^44^. EG011 treatment increased F-actin abundance as quantified by flow cytometry, confirming effective activation of the actin polymerization machinery (Figure 2H, S3A). Strikingly, this cytoskeletal restoration enhanced CTL-mediated clearance of HIV WT-infected cells (Figure 2I, S3B). These findings demonstrate that cytoskeletal disruption in HIV-infected cells is not merely correlative with CTL resistance but constitutes a mechanistically and pharmacologically reversible barrier to immune elimination.

### HIV Nef drives cytoskeletal disruption and contributes to CTL resistance

The observation that HIV-infected cells exhibit reduced stiffness even prior to CTL exposure suggests that viral components may directly contribute to cytoskeletal remodeling (Figure 2B, C). Among HIV-encoded proteins, the virulence factor Nef has been implicated in modulating actin dynamics and interfering with host cytoskeletal signaling pathways^45–48^.To determine whether Nef contributes to the cytoskeletal and mechanical phenotypes associated with CTL resistance, we first compared CD4^+^ T cells infected with WT or Nef-deficient (ΔNef) JRCSF HIV by confocal microscopy. WT-infected cells exhibited a marked reduction in cortical F-actin relative to uninfected controls (Figure 3A, B), and they also failed to spread normally on stimulatory anti-CD3/anti-CD28-coated surfaces (Figure 3A, C), indicative of impaired immunological synapse formation. This phenotype was largely reversed in ΔNef-infected cells, which displayed only a modest reduction in F-actin content and no spreading defect (Figure 3A-C). These results, which are consistent with prior reports^46–49^, indicate that Nef is the principal mediator of HIV-induced cytoskeletal disruption.

**Figure 3:**
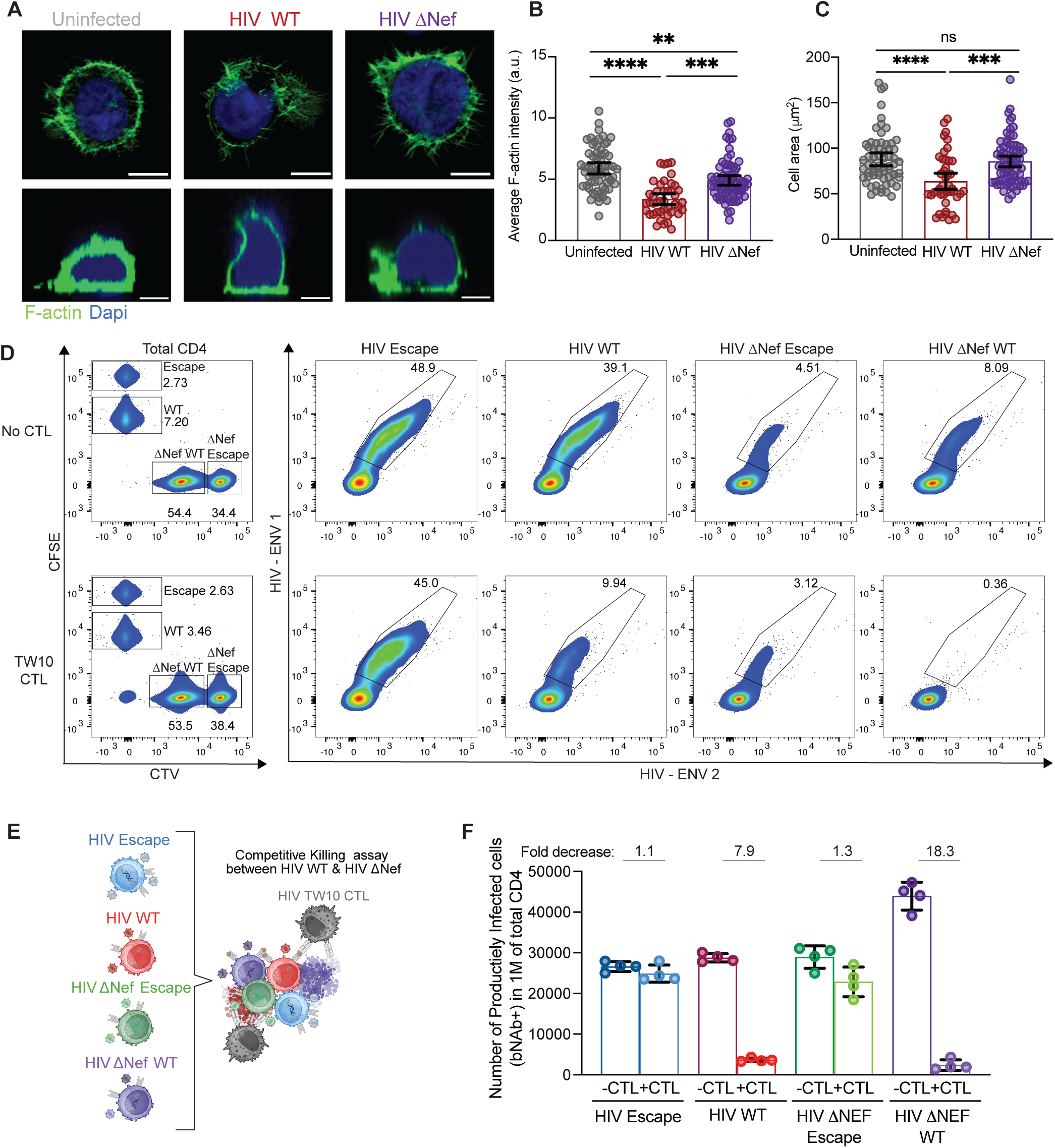
HIV Nef remodels the cytoskeleton to alter the mechanical proprieties of infected CD4^+^ T cells and enhance their resistance to CTL mediated killing. (A) Confocal imaging of F-actin (GFP) and cell spreading in uninfected (gray), HIV WT-infected (red), and HIV ΔNef-infected (purple) CD4^+^ T cells. Cell spreading area was measured on glass surfaces coated with poly-L-lysine and anti-CD3 antibodies. (B) Quantification of F-actin levels for the conditions described in (A). (C) Quantification of cell spreading for the conditions described in (A). Each point represents an individual cell; one-way ANOVA (p<0.0001). In (B) and (C), each point represents an individual cell; ** and **** denote p<0.01 and p<0.0001, respectively, calculated by one-way ANOVA. ns = not significant. (D) Flow-cytometry gating strategy for the competitive killing assay. To distinguish each infected population, cells were labeled with high or dim CFSE (HIV escape, HIV WT) or CTV (HIV ΔNEF escape, HIV ΔNef). (E) Schematic of competitive killing assay. HIV WT-, HIV escape-, HIV ΔNef- and HIV ΔNef escape-infected cells were co-cultured with or without TW10 CTLs for 16h at a ratio 1:1:1:1:2. (F) Absolute counts of HIV Env^+^ cells per 1M CD4^+^ T cells for each subpopulation described in (E) with or without CTL exposure. Fold decreases were calculated for each virus (- CTL vs. + CTL), mean ± 95% CI shown. Each point represents a technical replicate.

We next assessed whether Nef-dependent cytoskeletal remodeling contributes to the mechanical alterations induced by HIV. Unlike WT-infected cells, ΔNef-infected cells exhibited no measurable changes in cortical stiffness or membrane tension following coculture with CTLs, and their mechanical properties remained indistinguishable to those of uninfected cells (Figure S4A, B). These findings indicate that Nef expression is specifically required for the mechanical softening phenotype associated with CTL-resistant cells.

Building on the EG011 rescue data (Figure 2I), we reasoned that ΔNef-infected cells, which lack Nef-mediated mechanical changes, would be more vulnerable to CTL-mediated killing. To test this hypothesis, we performed a competitive killing assay in which WT- and ΔNef-infected CD4⁺ T cells, along with their corresponding TW10 epitope-escape counterparts, were co-cultured together in the presence or absence of TW10-specific CTLs (Figure 3D, E). This internally controlled design enabled simultaneous comparison of CTL sensitivity across infection types within the same well while minimizing variability. WT and Escape cells were mixed at a 1:1:1:1:2 ratio of WT:ΔNef:Escape:ΔNef-Escape:CTLs and incubated for 16 hours. ΔNef-infected cells were preferentially eliminated, exhibiting an 18.3-fold reduction compared to a 7.9-fold reduction for WT-infected cells, while both ΔNef-Escape and Escape populations were largely unaffected by CTL exposure (Figure 3E, F). These findings demonstrated that Nef confers a substantial survival advantage against CTL-mediated killing. They left unresolved, however, whether that advantage was driven by mechanical softening or by some other Nef dependent effect.

### Cytoskeletal remodeling via the Nef–PAK2 axis is as critical as MHC-I downregulation for CTL resistance

Nef is canonically recognized for its ability to promote CTL evasion through the downregulation of surface MHC-I (Figure 4A). However, our findings indicated that Nef also drives pronounced cytoskeletal dysregulation and cellular softening, a phenotype previously associated with impaired CTL-mediated killing in cancer models^32,34–36^. Among host pathways targeted by Nef, the PAK2-cofilin axis is one of the most conserved and mechanistically well-characterized^45,48–50^. Nef recruits and activates the serine/threonine kinase PAK2 (p21-activated kinase 2) within a multiprotein complex, leading to phosphorylation and inactivation of cofilin, an F-actin-severing protein essential for filament turnover and cytoskeletal dynamics (Figure 4A). While this pathway has been implicated in altered T cell polarization and migration, its role in modulating the vulnerability of HIV-infected cells to CTL-mediated killing has not been explored. Intriguingly, HIV-infected cells that survived CTL coculture displayed elevated levels of PAK2 transcription (Figure 1H).

**Figure 4:**
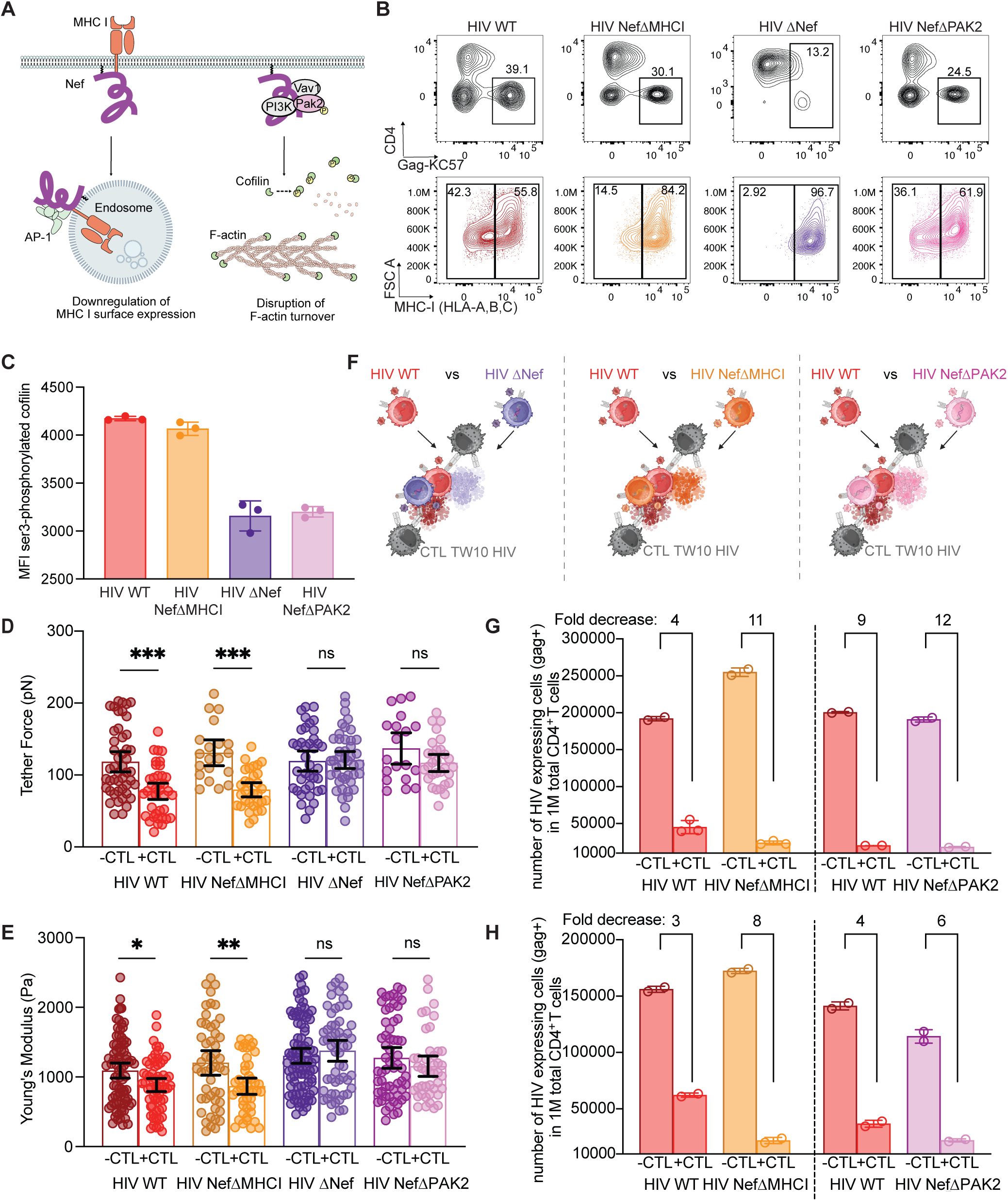
PAK2-dependent cytoskeletal remodeling by Nef confers CTL resistance comparable to MHC-I downregulation. (A) Schematic showing Nef’s interactions with AP-1 to downregulate MHC-I, and with PAK2 to phosphorylate and inhibit cofilin, thereby limiting F-actin turnover. (B) Flow cytometry plots of Gag^+^ cells showing CD4 and MHC-I downregulation with various mutants. (C) Mean fluorescence intensity (MFI) of Ser3-phosphorylated cofilin in HIV WT-, HIVΔNef-, HIV-NefΔMHCI-, or HIV-NefΔPAK2-infected CD4+ T cells. Data represents technical replicates with mean ± SD. (D and E) Membrane tether force (D) and cortical stiffness (E) of HIV WT-infected (red), HIV NefΔMHCI-infected (orange), HIV ΔNef-infected (purple) and NefΔPAK2-infected (pink) CD4^+^ T cells with and without CTL co-culture. Data are shown for one donor. *, **, and *** denote p≤0.05, p<0.01, and p<0.001, calculated by one-way ANOVA. Each point represents an individual cell; mean ± 95% CI from two independent experiments per condition. (F) Schematic of competitive killing assay in which equal numbers of HIV WT- and HIVΔNef-, HIV-NefΔMHCI-, or HIV-NefΔPAK2-infected cells were co-cultured with or without TW10 CTLs for 16h at a 1:1:1:1 ratio. (G and H) Competitive killing assays comparing HIV-WT with NefΔMHC-I or NefΔPAK2 variants in two donors. Absolute counts of Gag⁺ cells per 1M CD4⁺ T cells are shown with or without CTL exposure. Data represent technical replicates with mean ± SD.

To test whether Nef-mediated activation of PAK2 contributes to CTL resistance, and whether this effect is functionally distinct from that of MHC-I downregulation, we generated two Nef point mutants in the JRCSF background: one defective for MHC-I downregulation (M20A; NefΔMHC-I)^5^, and another bearing a hydrophobic patch mutation that abrogates PAK2 recruitment (L201A; NefΔPAK2)^51^ (Figure 4A, S4E). As expected, NefΔMHC-I HIV failed to induce MHC-I downregulation in infected CD4⁺ T cells, whereas NefΔPAK2 HIV suppressed surface MHC-I comparably to WT virus (Figure 4B, S4F). Conversely, phosphorylated cofilin levels were markedly reduced in NefΔPAK2-infected but not NefΔMHC-I-infected cells, consistent with loss of PAK2 signaling in the former but not the latter population (Figure 4C). Both mutants, however, maintained the ability to downregulate CD4, unlike ΔNef HIV (Figure 4B), confirming selective disruption of distinct Nef functions.

NefΔMHC-I-infected cells mirrored the mechanical profile of WT-infected cells, displaying reduced cortical stiffness relative to uninfected cells prior to CTL co-culture, with additional softening and membrane tension reduction amongst survivors (Figure 4D, E). In contrast, NefΔPAK2-infected cells phenocopied ΔNef-infected cells, maintaining cortical stiffness and membrane tension levels comparable to that of uninfected controls, with no significant differences even after CTL co-culture (Figure 4D, E). These results identify the PAK2-interacting domain of Nef as the principal mediator of mechanical remodeling and demonstrate that this function is genetically separable from Nef’s canonical role in MHC-I downregulation.

Finally, to assess how these distinct Nef functions influence CTL susceptibility, we performed competitive killing assays in which WT HIV-infected CD4^+^ T cells were co-cultured with either NefΔMHC-I- or NefΔPAK2-infected T cells in presence of HIV-specific CTLs (E:T 1:2, 1:1:1 overall ratio of WT:Nef mutant:CTLs) (Figure 4F). Both NefΔMHC-I- and NefΔPAK2-infected cells were markedly more susceptible to CTL-mediated clearance than WT-infected cells (Figure 4G, H). These results demonstrate that disruption of either MHC-I downregulation or PAK2-dependent cytoskeletal remodeling is sufficient to restore CTL-mediated clearance of HIV-infected cells. Strikingly, abrogation of the PAK2-interacting domain of Nef sensitized infected cells to CTL killing to an extent comparable to that observed upon the loss of MHC-I downregulation, establishing the PAK2-cofilin pathway as an equally potent and previously unrecognized mechanism of HIV immune evasion.

## Discussion

The present study establishes a previously unrecognized mechanical axis of immune evasion that operates in parallel with canonical biochemical mechanisms such as MHC-I downregulation. CTL-mediated killing is a mechanosensitive process: formation of a stable immune synapse, transmission of nanoscale forces, and perforin-dependent pore formation all rely on physical resistance from the target surface^29,32,33,39,40^. By diminishing cortical stiffness through cytoskeletal remodeling, HIV reduces these interfacial forces, weakening the delivery of lethal granules and diminishing killing efficiency^34,48^. Importantly, the extent of protection achieved through this mechanism is comparable to that of MHC-I downregulation, positioning mechanical modulation as a critical determinant of CTL susceptibility^5,37^.

Our results support a model of ‘collusion’ between cell-intrinsic mechanical properties of individual CD4^+^ T-cells and cytoskeletal remodeling induced by Nef. Cortical stiffness is inherently heterogeneous across T cells, reflecting stochastic variation in cytoskeletal organization and activation history. Nef exploits this diversity by activating the PAK2-cofilin pathway to suppress actin dynamics, further softening its host cell and amplifying the survival bias toward mechanically pliant targets. This represents a form of “selection through compliance,” in which HIV leverages cell-to-cell variation at the physical level to achieve the same evolutionary outcome that sequence variation achieves at the viral level: persistence under immune pressure. The cell-intrinsic contribution to mechanical resistance may help explain why certain infected T-cell clones disproportionately contribute to the long-lived reservoir^52–54^. Notably, the canonical MHC-I downregulation function of Nef also appears to be subject to ongoing selection on ART, with greater persistence associated with higher degrees of this activity^55^. Thus, the distinct properties of the provirus-host-cell chimera may evolve in composite fashion, with viral and cellular traits co-adapted to produce rare but exceptionally durable lineages of infected cells. These hybrid adaptations, genetic at the viral level and phenotypic at the cellular level, may together define the selective landscape that shapes the persistent reservoir.

It is tempting to speculate that Nef-induced mechanical softening evolved to complement the deficiencies of MHC-I downregulation as an evasion mechanism. Although cells lacking MHC-I are protected from CTLs, they are concomitantly sensitized to NK cells via “missing self” recognition. By contrast, cell softening is expected to attenuate both CTL and NK cell cytotoxicity by targeting the mechanosensitive properties shared by all cytotoxic lymphocytes^32,56,57^. This biomechanical mode of resistance is not without cost, however. Indeed, prior studies have demonstrated that Nef-PAK2 signaling impairs cell migration^51^, which could hinder viral dissemination, at least in the short term. That HIV sacrifices some of its capacity to spread in order to escape cytotoxic lymphocytes is a testament to the evolutionary imperative of immune evasion.

Transcriptomic profiling of CTL-resistant HIV-infected cells has revealed coordinated suppression of cytoskeletal remodeling genes and induction of apoptosis inhibitors, together with signatures of metabolic quiescence and reduced oxidative stress, features characteristic of long-lived, apoptosis-resistant memory T cells^21,37,38^. These convergent programs suggest that reservoir-harboring cells are selectively enriched across orthogonal axes of immune resistance; metabolic, biomechanical, and immunologic. This multifactorial architecture of evasion likely contributes to the persistent difficulty of clearing HIV reservoirs.

Our finding that the re-activation of actin polymerization with EG011 sensitized infected cells to CTLs demonstrates that mechanical resistance is not a fixed trait but a drug-modifiable property. That actin dynamics can be safely tuned in primary T cells suggests that the cytoskeleton may be a more tractable target than long assumed. That being said, global modulation of actin turnover would likely fall short of the specificity required for in vivo applications. Potentially more promising are interventions at the Nef–PAK2–cofilin interface, where viral specificity and host dependency converge. Small-molecule inhibitors that block Nef-PAK2 binding or allosteric modulators of newly defined Nef interaction sites could selectively reverse the cytoskeletal remodeling that underlies mechanical resistance while preserving essential F-actin functions. Building on this principle, mechanical re-sensitization could be developed as an orthogonal therapeutic axis that complements existing biochemical approaches. Agents that restore cytoskeletal integrity and target-cell rigidity could be combined with pro-apoptotic or epigenetic modulators, one reinstating the mechanical coupling required for cytotoxicity, the other enhancing antigen presentation and effector vigor.

Our results raise the possibility that other intracellular pathogens might also utilize cytoskeletal remodeling to biophysically evade the immune system. Microbes such as *Listeria monocytogenes*^58^, *Mycobacterium tuberculosis*^59^, and *Chlamydia trachomatis*^60^, manipulate the host cytoskeleton with remarkable spatial and temporal precision in order to support key stages of their lifecycle, including entry, replication, and egress. However, the consequences of these remodeling events for immune evasion remain unclear. Alongside canonical biochemically-based mechanisms, such as MHC downregulation, proteolytic degradation of antigenic peptides, and modification of microbial ligands, biophysical remodeling of host cells represents a parallel and potentially equally important resistance strategy.

Together, our findings define a mechanical dimension of immune evasion whereby HIV, through Nef-mediated cytoskeletal remodeling, transforms its host cell into a compliant target that resists cellular cytotoxicity. This strategy amplifies intrinsic cellular heterogeneity to ensure survival under immune pressure and potentially contributes to the emergence of long-lived reservoir lineages. Recognizing that immune escape can proceed through both biochemical and biophysical routes expands the conceptual framework of persistence and positions cytoskeletal mechanics as an actionable determinant for durable viral control and potential cure. By coupling molecular manipulation to biophysical adaptation, viruses achieve immune invisibility through force as well as form, broadening the known dimensions of host-pathogen coevolution.

## Acknowledgments

We thank all study participants who devoted time to our research, as well as the clinical research team involved in the study. This work would not have been possible without regents provided by the AIDS and Cancer Virus Program, Leidos Biomedical Research, Inc., Frederick National Laboratory for Cancer Research, supported with federal funds from the National Cancer Institute, National Institutes of Health, under contract HHSN261200800001E. We acknowledge the Biological Resources Branch Preclinical Repository, National Cancer Institute, for providing IL-2 and IL-15. We thank the molecular cytology core facility at MSKCC and the flow cytometry core at Weill Cornell Medicine for their assistance in this publication. This work was supported by the following NIH grants: R01MH130197, R01AI170239, R01AI087644, R37AI181626, R01AI176601, R01AI176943, R01AI170245, R01AI165031, UM1AI164562, UM1AI164565, R01AI150412, R21AI170246, R01AI147845, U01AI145921, R01AI167691, R21AI172554, and P30CA008748. The funders had no role in study design, data collection and analysis, decision to publish, or preparation of the manuscript. Figure and figure schematics were created using Flow Jo, ImageJ, GraphPad Prism, BioRender.com, Inkscape, Cytoscape and Adobe Illustrator.

## Author contributions

L.L, F.M, M.H, and R.B.J conceived the study. L.L and F.M designed and conducted experiments, and interpreted the results A.H, E.L, J.W, and C. S assisted in data compilation E.H, A.H, P.Z, and D.B performed bioinformatic analyses. K.L.C, P.S, and E.N contributed essential reagents or analytical tools. C.K provided clinical samples or patient data. R.B.J, and M.H supervised research. F.M, L.L, M.H, and R.B.J wrote the manuscript with input from all authors. All authors reviewed, edited, and approved the final manuscript.

## Methods

### HIV infection model in cultured central memory CD4+ T cells (Tcm)

HIV-infected CD4^+^ T cells were generated using a previously described HIV latency model^61^. Briefly, peripheral blood mononuclear cells (PBMCs) were isolated from leukapheresis samples obtained from HIV negative donors (Stemcell technologies). Naïve CD4^+^ T cells were purified from thawed PBMCs using the EasySep™ Human Naïve CD4^+^ T Cell Isolation Kit. Purified naïve CD4^+^ T cells were activated with human anti-CD3/CD28-coated magnetic beads in the presence of human anti-IL-4 (1µg/10^6^ cells), human anti-IL-12 (2µg/10^6^ cells), and Tumor Growth Factor (TGF)-β1 (10ng/10^6^ cells) to prevent cell polarization. Cells were maintained at a density of 1x10^6^ cells/mL in culture media supplemented with 50 IU/mL recombinant human IL-2. On day 7, cultured Tcm cells were infected with one of several HIV JRCSF variants: wild-type JRCSF (WT HIV), TW10escape JRCSF (Escape HIV), ΔNef JRCSF (HIVΔNef), JRCSF Nef M20A mutant (HIV NefΔMHC-I), JRCSF Nef L201A mutant (HIV NefΔPAK2) or TW10escape ΔNef JRCSF (Escape HIVΔNef). One-quarter of the cells were infected by spinoculation with virus at 1000 × g for 2h at 37°C. The infected cells were then mixed with the remaining three-quarters of uninfected cells and cultured at 2×10^6^ cells/mL in R10-50 media. On day 10, cells were crowded into 96-well round-bottom plates at a density of 2×10^5^s in 200 µL to enhance viral transmission. On day 13, cells were cryopreserved in FBS with 10% DMSO and stored in liquid nitrogen for future use.

### Virus production and infections

The HIV JRCSF wild-type (WT HIV) plasmid was obtained from the NIH AIDS Research & Reference Reagent Program. The HIV JRCSF TW10 escape (Escape HIV), HIV JRCSF ΔNef, HIV JRCSF NefΔPAK2, and HIV JRCSF NefΔMHC-I plasmids were synthesized by GenScript. The HIV JRCSFΔNef plasmid was generated from the WT plasmid by a 20 amino acid deletion following serine 56, resulting in a premature stop codon due to a frameshift. The HIV JRCSF NefΔPAK2 mutant was generated by substituting leucine 201 with alanine, and the HIV JRCSF NefΔMHC-I mutant was generated by substitution of methionine 20 with alanine. Plasmids were expanded in E.coli Stlb3 competent cells and purified using the QIAGEN Plasmid Plus Maxi Kit. Viral stocks were produced by transfecting full-length plasmids into HEK293T cells using FuGENE 6 Transfection Reagent in Optimem. At 24h post-transfection, the culture media was replaced with DMEM with 10% FBS. After an additional 48h, viral supernatants were harvested, filtered through 0.45µm filters, and concentrated overnight at 4°C using the PEG-it virus precipitation solution. Concentrated viruses were aliquoted and stored at −80°C until use. For infection experiments, cells were exposed to virus at a 1:5 cell-to-virion ratio, which yielded optimal infection efficiency and cell viability.

### Bystander and survivor CTL Killing assays

HIV WT and HIV escape infected cells generated as described above were thawed and rested for 2 days at 2 × 10^6^ cells/mL in media supplemented with 50 IU/mL IL-2. Prior to coculture, HIV expressing cells were enriched by depleting cells with high surface CD4 expression using the EasySep™ Human CD4 Positive Selection Kit II. WT HIV infected cells were labeled with CFSE (0.2 µM), and Escape HIV infected cells were labeled with CTV (0.5 µM). CFSE labeled WT HIV infected cells and CTV labeled escape HIV infected cells were mixed at a 2:1 ratio. The mixed target cells were then cocultured with or without TW10-specific CTL clones recognizing the HIV Gag epitope TW10 (TSTLQEQIGW). CTLs were labeled with CTB (0.5 µM) and added at an E:T ratio of 1:1. Anti-CD107a APC-cy7 (1:200) was included in the culture to monitor CTL degranulation. Killing assays were performed in 96 wells round bottom plates in culture media supplemented with IL-2 (50 IU/mL) and IL-15 (0.51 ng/mL) for 16h at 37°C. Following coculture, cells were stained for live/dead, surface CD3 and CD4, and for HIV using either extracellular or intracellular staining. Extracellular staining was performed with HIV Env specific antibodies, PGT121 (Env 1) and trimer N6/PGDM1400x10E8 (Env 2), conjugated with the Zenon labeling kit (1 ug per million of cells). Intracellular HIV Gag staining was performed with the KC57-RD1 antibody (1:500). For sequencing and for membrane tension and cortical stiffness measurements, cells were sorted based on CFSE or CTV fluorescence and double positivity for HIV Env antibodies. Intracellular Gag staining was used for flow cytometric analysis on FlowJo software.

### Competitive killing assay

Competitive killing assays were designed to evaluate CTL-mediated elimination of distinct HIV-infected target cell populations within a mixed culture. In this assay, CD4+ T cells infected with different HIV variants were cocultured with effector CTLs. For the first set of experiments, competition assays were performed between WT HIV and ΔNef HIV-infected cells, each paired with their corresponding escape virus-infected counterparts. WT and escape HIV-infected cells that downregulated CD4 were first enriched using the EasySep™ Human CD4 Positive Selection Kit II. WT HIV and escape HIV-infected cells were labeled with low (0.5 µM) and high (5 µM) CFSE levels, respectively, while ΔNef HIV and ΔNef escape HIV-infected cells were labeled with low (0.5µM) and high (5µM) CTV levels, respectively. The absolute numbers of infected cells were normalized across samples based on intracellular HIV p24 staining. Equal numbers of each labeled p24+ infected population were then mixed and cocultured with or without HIV-TW10 CTLs at a 1:1 effector-to-recognized-target ratio (where recognized targets were WT HIV- and ΔNef HIV-infected cells). Cocultures were incubated for 16 hours at 37°C in 96-well-round-bottom plates (maximum of 1×10^6^ total cells per well) in complete RPMI media supplemented with 50 IU/mL IL-2 and 0.5 ng/mL IL-15. In a separate set of competitive killing assays, WT HIV-infected cells were cocultured with either ΔNef HIV-, NefΔPAK2 HIV-, or NefΔMHC-I HIV-infected cells. Each infected population was labeled with either CFSE (0.5 µM) or CTV (0.5 µM) and normalized for equivalent frequencies of infected expressing cells (p24+) before coculture. These mixed targets were incubated with or without CTLs for 16 hours under the same culture conditions as above. Following incubation, cells were harvested and stained for CD3, CD4, CD8, Live/Dead and HLA-A/B/C. The frequency and total number of infected cells were determined by either extracellular Env staining (using PGT121 and trimer N6/PGDM1400x10E8 antibodies conjugated with a Zenon labeling kit) for microscopy experiments or intracellular p24 Gag staining (clone KC57) for infected cells survival analysis by Flow cytometry. All the data were analyzed with FlowJo software.

### RNA-seq data analysis

Raw reads were quality checked with Fast QC v0.11.7 (http://www.bioinformatics.babraham.ac.uk/projects/fastqc/). Reads were aligned to the human reference genome (GRCh38.p12) using STAR v2.6.0c^62^ with default parameters. Gene abundances were calculated with featureCounts v1.6.2 ^63^ using composite gene models from Gencode release 28 ^64^. Principle component analysis was performed using the plotPCA function from DESeq2 v1.32.0 ^65^. Differentially expressed genes were determined with DESeq2 v1.32.0 using Wald tests (q < 0.05). Gene set enrichment analysis was performed using fgsea v1.18.0 ^66^; genes were ordered by the DESeq2 Wald statistic. Gene sets were retrieved from the Broad Institute’s MSigDB collections^67^. Only pathways with an *P-*adj value < 0.05 were considered enriched. Over-representation testing of gene ontology terms (GO) was performed using clusterProfiler v4.0.5 (q < 0.05) ^68^. Expression heatmaps were generated using variance-stabilized data, with the values centered and scaled by row.

### Fixed confocal microscopy and quantification

Glass-bottom petri dishes (FD35-100, World Precision Instruments) were plasma treated for 2 min on high setting and then coated with poly-l-lysine (Sigma) for 1 h at room temperature. Dishes were then washed and treated with 5 μg/mL of anti-CD3 (OKT3, 14-0037-82, Thermo Fisher) overnight at 4⁰C. After thoroughly washing with PBS, cells were plated for 1 h at 37⁰C and then fixed with 2% paraformaldehyde (PFA, Electron Microscopy Sciences) for 10 min. The cell membrane was permeabilized with cold, 0.1 % Triton X-100 solution for 5 min, and the samples were blocked with 10% BSA for 1 h. Staining was performed using Alexa488-conjugated phalloidin (1:1000, A12379, Thermo Fisher), to visualize actin, and DAPI (1:1000, D1306, Thermo Fisher), to visualize cell nuclei, for 2 h at room temperature. Samples were imaged using a Leica Stellaris 8 point scanning confocal microscope fitted with a 63×/1.4 NA oil objective. Image stacks encompassing entire cells were acquired using 0.3-µm z-sectioning. Image analysis was performed using open-source Fiji ImageJ software (v 2.9.0). All fluorescent images for a given channel were recorded using the same imaging parameters (laser, intensity, exposure time, and threshold). For analysis, all image stacks were subjected to intensity thresholding to establish the space occupied by cells, after which the sum intensity of Alexa488-conjugated phalloidin within the cellular volume was determined.

### Membrane tension measurements

Plasma membrane tension was quantified using C-Trap optical tweezers (Lumicks BV, Netherlands). An IR laser beam (50 mW, 1064 nm) was tightly focused through a series of mirrors, beam expanders and a high numerical aperture objective lens (63×/1.2 NA, Nikon Instruments) to form a steerable optical trap. Cells were seeded for 1 hour on glass-bottom petri dishes coated with 5 μg/mL of anti-CD3. To measure plasma membrane tension, polystyrene beads (2.2 μm, Spherotech Inc, IL) were coated with concanavalin A (50 μg/mL, Thermo/Sigma) and added to the cell culture medium in the dish. Beads were momentarily placed in contact with the cell membrane, and tethers were then extruded by moving the bead away from the cell perpendicularly at a speed of 2 μm/s. Force measurements were made using the Lumicks Bluelake software suite by capturing the exiting trapping light with a high numerical aperture condenser lens (63×/1.45, oil immersion, Zeiss AB, Germany) and measuring bead displacement in the trap with position-sensitive detectors through back focal plane interferometry. Membrane tether breaking was documented as a sharp discontinuity in tether force during tether extrusion, with breaking distance measured from simultaneously collected brightfield images using FIJI. Data analysis was performed using Python 3.8.0.

### Young’s modulus measurements

Cells were seeded on glass-bottom petri dishes coated with 5 μg/mL of anti-CD3. After 1 h, the cells were fixed for 10 min with 2% PFA and then kept in complete RPMI medium during the acquisition of single force measurements. Experiments were performed at room temperature with the Asylum Research MFP-3D-BIO AFM microscope (Oxford Instruments) using cantilevers with a 1 μM diameter spherical probe (SAA-SPH-1UM, nominal spring constant k = 0.25 N/m, Bruker). Before each experiment, the exact spring constant of the cantilever was determined using the thermal noise method and its optical sensitivity determined using a PBS-filled glass bottom petri dish as an infinitely stiff surface. Between 40 to 70 cells were measured for each experimental group during each session. At least 3 measurements were collected per cell. The maximal force applied was 500 pN, leading to indentation depths on the order of 1 μM. Force curves were fitted according to the Hertz model (Igor Pro, Wavemetrics). Data fitting was performed in the range from 0 to 50% of the maximum applied force to consider only measurements within the first 1 μM of indentation. The following settings were used: tip Poisson vtip = 0.19, tip Young’s modulus Etip = 68 GPa, and sample Poisson vsample = 0.45.

### EG011 treatment

The small molecule EG011^45^, a WASP activator, was obtained in powder form and dissolved in DMSO at a concentration of 10 mM. The solution was aliquoted into single use 10 μL vial and stored at -20°C until used. EG011 was optimized for use at a final concentration of 1.5 µM in the culture medium, resulting in a final DMSO concentration of 0.01%. DMSO was used a vehicle control in all the experiments. For killing assays with HIV infected cells, the infected cells were pretreated with EG011 or vehicle for 6 h before co-culture with or without CTLs. The coculture was maintained in media containing EG011 or vehicle for 16h.

### Phospho-flow staining

Phospho-flow cytometry was performed to assess intercellular signaling of phosphorylated cofilin (Ser3) in cells infected with WT HIV, ΔNef HIV, NefΔPAK2 HIV, or NefΔMHC-I HIV. Infected cells were thawed and rested overnight, after which and 1.5×10^5^ were aliquoted per well into 96-well round bottom plates. Triplicate samples were prepared for each virus condition. Cells were first stained for surface markers in FACS buffer (PBS containing 2% FBS and 2mM EDTA) with Live/Dead aqua viability dye and CD4 antibody for 20 minutes at 37°C. After staining, cells were fixed with pre-warmed 2% PFA (Electron Microscopy Sciences) for 10 minutes at 37°C. After washing, fixed cells were permeabilized by the gradual addition of ice-cold BD Phosflow™ Perm buffer III and then incubated on ice for 30 minutes. Following permeabilization, cells were washed with FACS buffer and stained for intracellular markers, including anti-phospho-cofilin (Ser3) antibody (1:500 dilution) and anti-p24 Gag (clone KC57) for 30 minutes at room temperature. After washing, samples were incubated with AF 594-conjugated anti-rabbit secondary antibody (1:500 dilution) for 30 minutes at room temperature. Finally, cells were washed twice with FACS buffer and resuspended in 200 μL of FACS buffer for acquisition. Data were analyzed using FlowJo software.

---

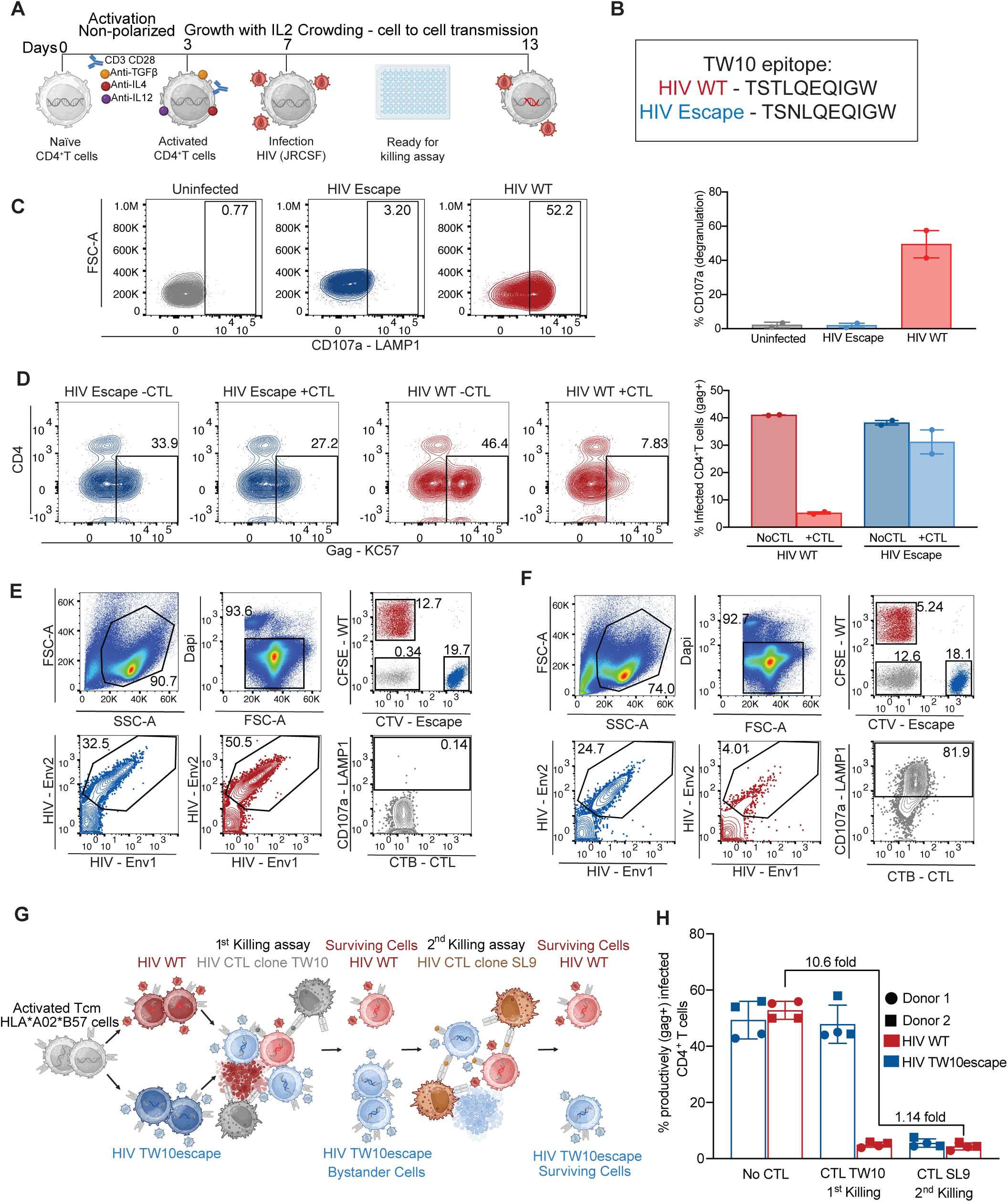

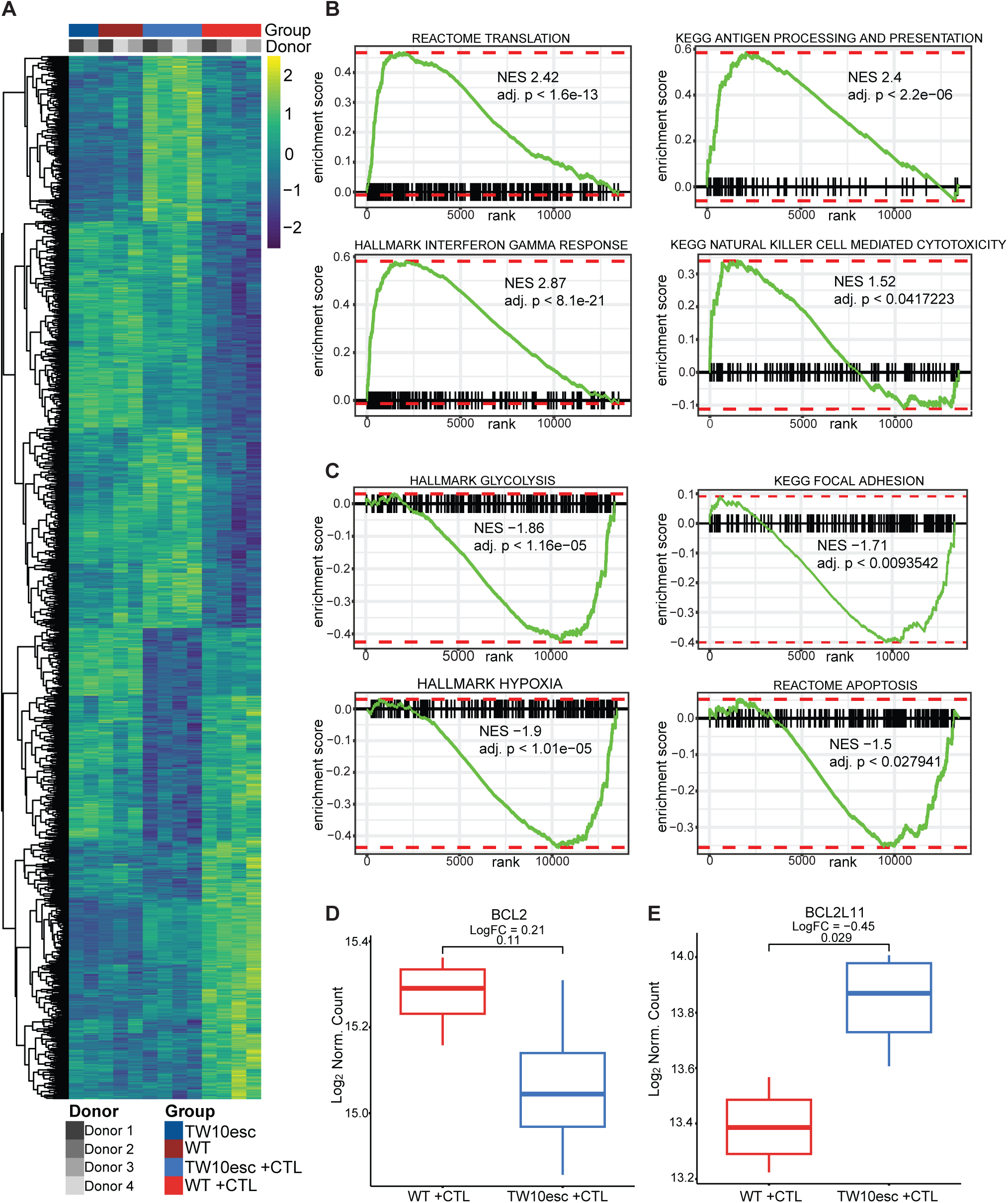

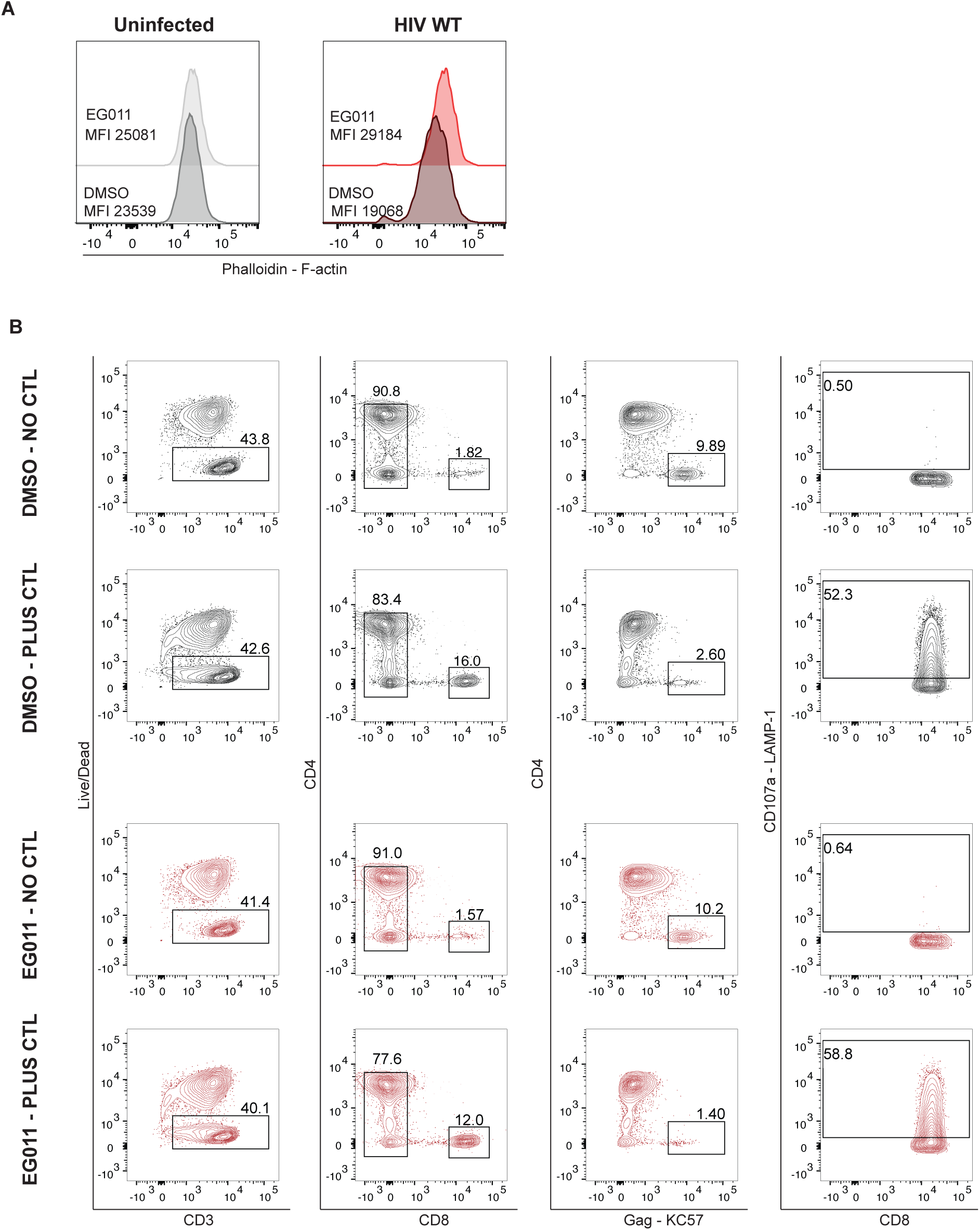

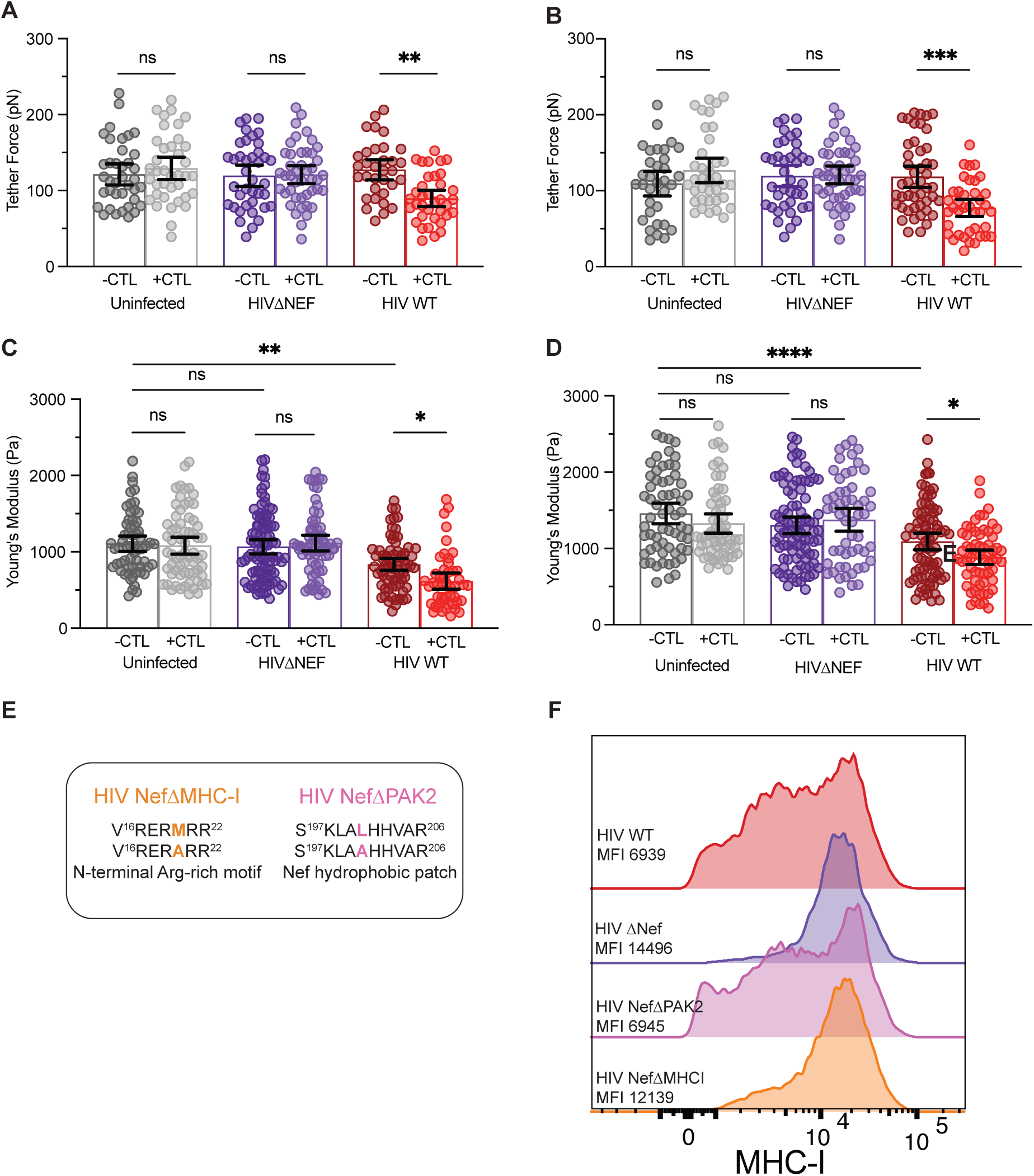

