## Supplementary Figure Legends for "HIV Nef amplifies mechanical heterogeneity to promote immune evasion"

**Supplemental figure legends:**

**Figure S1: Characterization of HIV infection model and CTL recognition of WT and TW10 escape variants.** (A) Schematic of the HIV infection model in primary activated central memory $CD4^+$ T cells. (B) Single-amino acid mutation in the TW10 epitope of HIV JRCSF converting threonine (T) 242 in *gag* to asparagine (N), preventing TW10 peptide binding to MHC-I. (C) Left, representative flow cytometry plots showing LAMP-1 (CD107a) staining of TW10 CTLs after coculture with uninfected, HIV WT-infected, or HIV TW10 escape-infected CD4+ T cells. Right, quantification of $CD107a^+$ CTLs under each condition. (D) Flow cytometry gating (left) and percentage of $Gag^+$ T cells (right) in HIV WT and HIV escape infected cells co-cultured with or without TW10 CTLs. (E, F) Flow cytometry gating used to sort HIV-WT and HIV-escape-infected cells for bulk RNA sequencing. HIV-WT- and HIV escape-infected cells were labeled with CFSE and CTV, respectively, and co-cultured in the presence (H) or absence (G) of TW10 CTLs for 16 h. Infected cells were identified and sorted based on co-staining with HIV-Env specific antibodies. (G) Schematic of the sequential killing assay involving two rounds of CTL-mediated killing using clones specific for the Gag-TW10 or Gag-SL9 epitopes. (H) Percentage of productively infected (Gag+) CD4+ T cells during sequential killing assays. In the first round, TW10-specific CTLs were co-cultured with HIV WT- (red) and HIV escape- (blue) infected cells. In the second round, remaining HIV WT and HIV escape infected cells were co-cultured with SL9-specific CTLs. Fold changes in infected (CD4+ HIV-Gag+) cells before and after killing are shown for two donors in technical duplicates

**Figure S2: Multiple intrinsic pathways contribute to HIV-infected cell resistance to CTL-mediated killing**. (A) Heatmap of 2,243 differentially expressed genes (DEGs) identified between HIV-WT and HIV-escape with or without CTL coculture (row-normalized log counts per million; Wald test - adjusted p-value < 0.001). (B-C) Gene set enrichment analysis (GSEA) of pathways upregulated (B) or downregulated (C) in survivor cells, featuring Reactome, Hallmark, and KEGG gene sets.  (D-E) Boxplot showing log2 fold change in normalized expression counts of BCL2 (D) and BCL2L11 (BIM) (E) between survivor (HIV WT) and bystander (HIV escape) populations. Statistical significance was determined by Wilcoxon test.

**Figure S3: Flow cytometry gating and analysis of EG011 treatment.** (A) MFI of F-actin in uninfected and HIV WT infected cells treated with EG011 (1.5 $\mu$M) or vehicle (DMSO) for 24 h. (B) Flow cytometry gating strategy for the killing assay of HIV WT-infected cells pretreated for 6 h with EG011 (1.5$\mu$M) or vehicle (DMSO), followed by co-culture with or without TW10 CTLs in the presence of EG011 (1.5$\mu$M) or vehicle for 16 h.

**Figure S4: HIV ΔNEF does not alter the cortical stiffness or membrane tension of infected cells.** (A-B) Tether force (indicative of membrane tension, pN) of uninfected (gray), HIV WT-infected (red) and HIV ΔNEF-infected (purple) $CD4^+$ T cells before and after CTL-mediated killing. Data are shown for two donors (donor 1: (A), donor 2: (B)). (C-D) Cortical stiffness (Young's modulus, Pa) of uninfected (gray), HIV WT-infected (red) and HIV ΔNEF-infected (purple) CD4+ T cells before and after CTL-mediated killing. Data are shown for two donors (donor 1: C, donor 2: D). In A-D, *, **, ***, and **** denote $p \leq 0.05$, $p<0.01$, $p<0.001$, and $p<0.0001$, calculated by one-way ANOVA. Each point represents an individual cell; mean ± 95% CI from two independent experiments per condition. (E) Diagram of single-amino acid mutagenesis in JRCSF Nef disrupting MHC-I downregulation (M20A; NefΔMHC-I) or PAK2 interaction (L201A; NefΔPAK2). (F) Representative histograms showing HLA-A, B, C (MHC-I) staining of CD4+ T cells infected with HIV WT-, HIVΔNef-, HIV-NefΔMHC-I, or HIV-NefΔPAK2.
